# Stability and expression of SARS-CoV-2 spike-protein mutations

**DOI:** 10.1101/2022.03.21.485157

**Authors:** Kristoffer T. Bæk, Rukmankesh Mehra, Kasper P. Kepp

**Affiliations:** DTU Chemistry, Technical University of Denmark, Building 206, 2800, Kongens Lyngby, Denmark.; Department of Chemistry, Indian Institute of Technology Bhilai, Sejbahar, Raipur-492015, Chhattisgarh, India.

**Keywords:** SARS-CoV-2, covid-19, spike protein, mutations, protein stability

## Abstract

Protein fold stability likely plays a role in SARS-CoV-2 S-protein evolution, together with ACE2 binding and antibody evasion. While few thermodynamic stability data are available for S-protein mutants, many systematic experimental data exist for their expression. In this paper, we explore whether such expression levels relate to the thermodynamic stability of the mutants. We studied mutation-induced SARS-CoV-2 S-protein fold stability, as computed by three very distinct methods and eight different protein structures to account for method- and structure-dependencies. For all methods and structures used (24 comparisons), computed stability changes correlate significantly (99% confidence level) with experimental yeast expression from the literature, such that higher expression is associated with relatively higher fold stability. Also significant, albeit weaker, correlations were seen for ACE2 binding. The effect of thermodynamic fold stability may be direct or a correlate of amino acid or site properties, notably the solvent exposure of the site. Correlation between computed stability and experimental expression and ACE2 binding suggests that functional properties of the SARS-CoV-2 S-protein mutant space are largely determined by a few simple features, due to underlying correlations. Our study lends promise to the development of computational tools that may ideally aid in understanding and predicting SARS-CoV-2 S-protein evolution.

## Introduction

The pandemic caused by severe acute respiratory syndrome coronavirus 2 (SARS-CoV-2)[1–3] has led to intensive research into its spike-protein (S-protein, **Figure 1a**), whose evolution may lead to changes in its surface that reduce the efficiency of vaccine-induced antibodies.[4–6] The S-protein is a glycosylated homo-trimer on the surface of coronaviruses responsible for their characteristic shape.[7, 8] During entry of the virus particle into human host cells, the S-protein binds to the cell-surface receptor angiotensin-converting enzyme 2 (ACE2), **Figure 1b**.[9–11]

**Figure 1:**
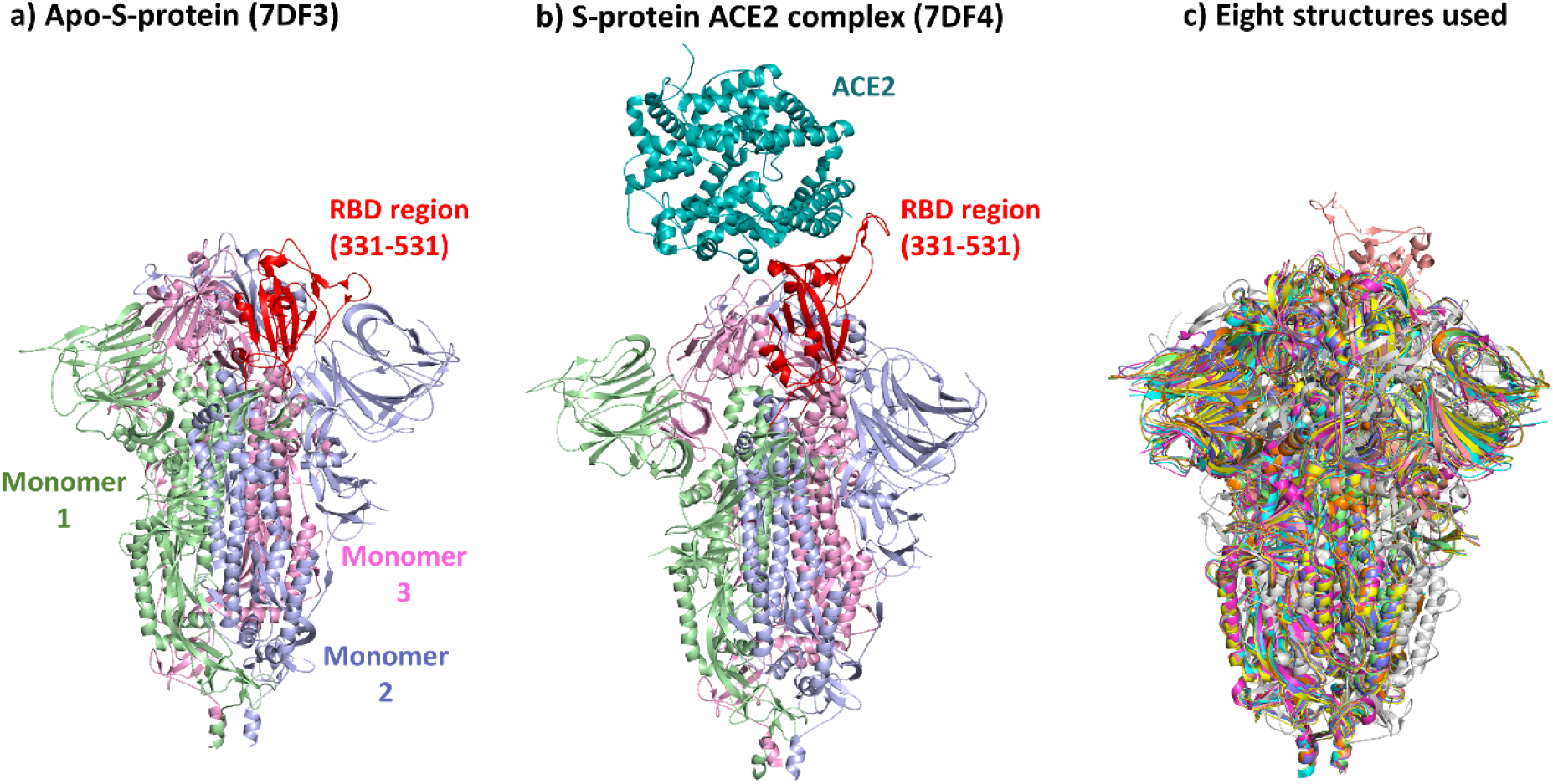
Structure of the SARS-CoV-2 S-protein, studied in this work. **(a)** Representative structure of the apo-S-protein state (PDB: 7DF3)[38] with the studied mutated sites of the RBD shown in red color. **(b)** Representative cryo-EM structure of the S-protein in complex with ACE2, (PDB: 7DF4). **(c)** Overlay of the eight apo-S-protein structures used in this study to account for structural heterogeneity when computing mutation effects.

Due to the importance of S-protein epitopes for immune recognition, presentation of the S-protein is the rationale behind a range of important vaccines.[12, 13] The presence of prominent antibodies in a population may lead to selection pressure to change the S-protein surface to evade the antibodies learned from vaccination or infection with earlier variants.[14] Such antigenic drift may lead to variants capable of escaping vaccine-induced immunity, a problem that is likely to persist for many years. The omicron variant has been a hallmark example of such evolution, leading to a very large number of breakthrough infections in late 2021 and early 2022.[15]

Understanding such evolution effects requires protein structural data.[16–19] Since function is structure-dependent, the structure is the platform on which the amino acid evolution occurs, with evolution rates typically depending on the structural context of the amino acid site.[20–22] The recent technical breakthroughs in cryo-electron (cryo-EM) microscopy of large molecules such as proteins[23–27] are an excellent example of basic science providing essential innovation enormous utility, enabling publication of hundreds of SARS-CoV-2 S-protein structures during the last two years, both of the protein alone in the apo-state (apo-S-protein), in various conformation states, and in complex with a large range of antibodies and ACE2 at resolutions typically at 2-4 Å.[28]

Among the selection pressures acting on a protein beyond its direct function is the need to maintain the overall fold stability and translational fidelity, and thus evolution of new mutations often occurs with a tradeoff not to undermine these properties.[17, 29–32] Before fusion with a host cell, the S-protein is in a metastable conformation state under selection to evade antibodies and enhance ACE2 binding, which requires appropriate conversion to a conformation state with the receptor binding domain (RBD) in an up-wards conformation state.[33–37]

Understanding the importance of fold stability-function tradeoffs within the S-protein could be important for estimating the tolerability of new mutations and the likelihood and fitness of new emerging variants, which will remain an important topic even in the post-pandemic period.[39] To understand this, we can compute the stability effects of the many mutations possible in the RBD. Many programs exist based on machine-learning or energy-based force-fields that can compute changes in protein stability upon mutation,[21, 40–47] with variable accuracy and issues relating to systematic errors (biases) and results being dependent on input structures used.[42, 48–52] The few experimental S-protein stability data from time-consuming differential scanning calorimetry or denaturation experiments prevent statistically meaningful analysis. Instead, we hypothesized that expression levels, which are available for very many SARS-CoV-2 S-protein mutations, may be a proxy of fold stability, as argued previously[53] and explored further below.

## Methods

### Data for expression and ACE2 binding

Data for the effect of S-protein RBD single-point mutations on yeast expression and ACE2-binding were published by Starr *et al*. 53] These mutation sites are marked in red color in **Figure 1a**. The effect of mutations on expression is reported as the difference in log-mean fluorescence intensity (MFI) relative to wild-type (ΔlogMFI = logMFI_variant_ – logMFI_wild-type_), such that a positive value indicates higher RBD expression.[53] The effect of mutations on ACE2-binding (**Figure 1b**) is calculated from the apparent dissociation constants (*K_D, app_*) and shown as the difference in log10(*K_D,app_*) relative to wild-type (Δlog10(*K_D,app_*) = log10(*K_D,app_*)_wild-type_ – log10(*K_D,app_*)_variant_), such that a positive value indicates higher variant ACE2 binding.[53] There were two independent measurements (one from each of two independent libraries) for each mutation, providing an important way to assess the quality and reproducibility of the individual data points.

### Data curation

There is an overall good correlation between the results obtained from the two independent libraries (**Figure S1a**). We removed outliers by filtering out observations where the two replicates are different (for binding *or* expression), defined as residuals > 1, or where data for one replicate is missing. The rationale is that data points not similar in the two experimental replicates cannot be considered reproduced and may erroneously affect analysis. We furthermore removed data points with effects on binding in either replicate < −4.5 to avoid values near the detection limit (see **Figure S1a**). The correlation between the replicates after curation is shown in **Figure S1b**. Removing outliers and data points at the detection limit also excluded stop codon mutations (**Figure S2**). The described curation removed 685 of the original 4221 data points. Further analyses were performed using the remaining 3536 data points and the average of the binding and expression data from the two independent libraries, with detailed data collected in the supplementary file **Table_S1.csv**.

### Structures and computer models used to compute stability effects

For each mutation in the dataset, the change in protein free energy of folding (ΔΔG, kcal/mol) was computed using three different methods: DeepDDG[54], mCSM Protein Stability[45] (mCSM), and SimBa-IB[55]. DeepDDG and mCSM were accessed via their respective web servers (http://protein.org.cn/ddg.html, http://biosig.unimelb.edu.au/mcsm/stability), and Simba-IB was run from the command-line (http://github.com/kasperplaneta/SimBa2).

We have previously shown that analysis of functional properties can depend on input structure used,[56] and such heterogeneity is also seen in published cryo-EM structures of the S-protein.[28] Accordingly, we used eight experimental cryo-EM structures of the apo-S protein from the Protein Data Bank (PDB): 6VXX,[35] 6X6P,[57] 6X79,[58] 6Z97,[59] 6ZB4,[60] 7CAB,[61] 7DDD,[62] and 7DF3,[38] to account for such heterogeneity. These structures are shown in structural overlay in **Figure 1c**.

The relative solvent accessible surface area (RSA) of the mutated sites were calculated using SimBa-IB[55], which uses FreeSASA for this task.[63] Because the eight PDB structures represent homo-trimers, the ΔΔG and RSA values reported in this study are average ΔΔG and RSA values for the three chains (A, B, and C) of each structure.

It is important to note that the cryo-EM structures discussed may not reflect very precisely the real conformations of the S-protein at physiological temperature (37 °C): Cryo-EM structures are typically obtained with samples deposited with vitrified ice and rapidly cooled using a cryo-agent.[24, 25, 64] The freezing may remove some conformational dynamics.[65–69] Conformational changes of the S-protein may also be temperature-dependent.[70] In addition, the physiologically relevant state of the S-protein tends to be heavily glycosylated, which none of the experimental assays studying mutation impacts so far has addressed. Our goal is to identify whether experimental apo-S-protein data can be approximated by computed data, without translating these to the physiological significance of these data.

## Results and discussion

### Experimental and computed genotype-phenotype heat maps

Our main interest was to investigate whether experimentally measured expression and ACE2 binding of S-protein RBD mutants[53] can be related to the thermodynamic stability of the mutants. Amino acid substitutions often change the stability of a protein with a tendency for the distribution of such effects to be skewed towards destabilization.[71] Mutations in RBD affect expression of the S-protein and binding to ACE2 differently, but as discussed before[53] there is a relatively strong correlation (R=0.59) between the effect of mutations on expression and ACE2 binding (**Figure S3a**). This relationship is not intuitive but could relate to underlying effects such as ACE2 recognition and stability both relating to the mutating site’s solvent accessibility, or correlations between codon use[72] which affects replication efficiency[73] and amino acid properties. Expression levels could also affect apparent binding constants even at the same specific ACE2 affinity, even if this is apparently accounted for, due to complex (e.g., tertiary) interactions. The large number of mutations having a small or “nearly neutral” impact, as noted by the authors[53] and reasonably expected, may also spuriously affect the relationships. However, if the data are grouped to adjust for the skewed distribution, the correlation between expression and binding becomes even larger and quite remarkable (R=0.88) (**Figure S3b**).

To assess the correlation between the effects of mutations on expression, ACE2 binding and protein stability, we computationally predicted the stability changes (ΔΔG) caused by the mutations in RBD using three state-of-the-art methods (DeepDDG, mCSM, and SimBa-IB) and compared with the experimentally observed changes in expression and binding. **Figure 2a** and **Figure 2b** show heat maps build from the experimental data by Starr et al.[53] after curation for non-reproducible replicates as described in Methods. These experimental heat maps are compared with the computational heat maps of ΔΔG derived in the present work, using DeepDDG (**Figure 2c**), mCSM (**Figure 2d**), and SimBa-IB (**Figure 2e**). SimBa produces more stabilizing trends overall, as it was developed to handle destabilization biases (the method performs similarly to other methods in benchmarks despite this feature)[55, 74].

**Figure 2.**
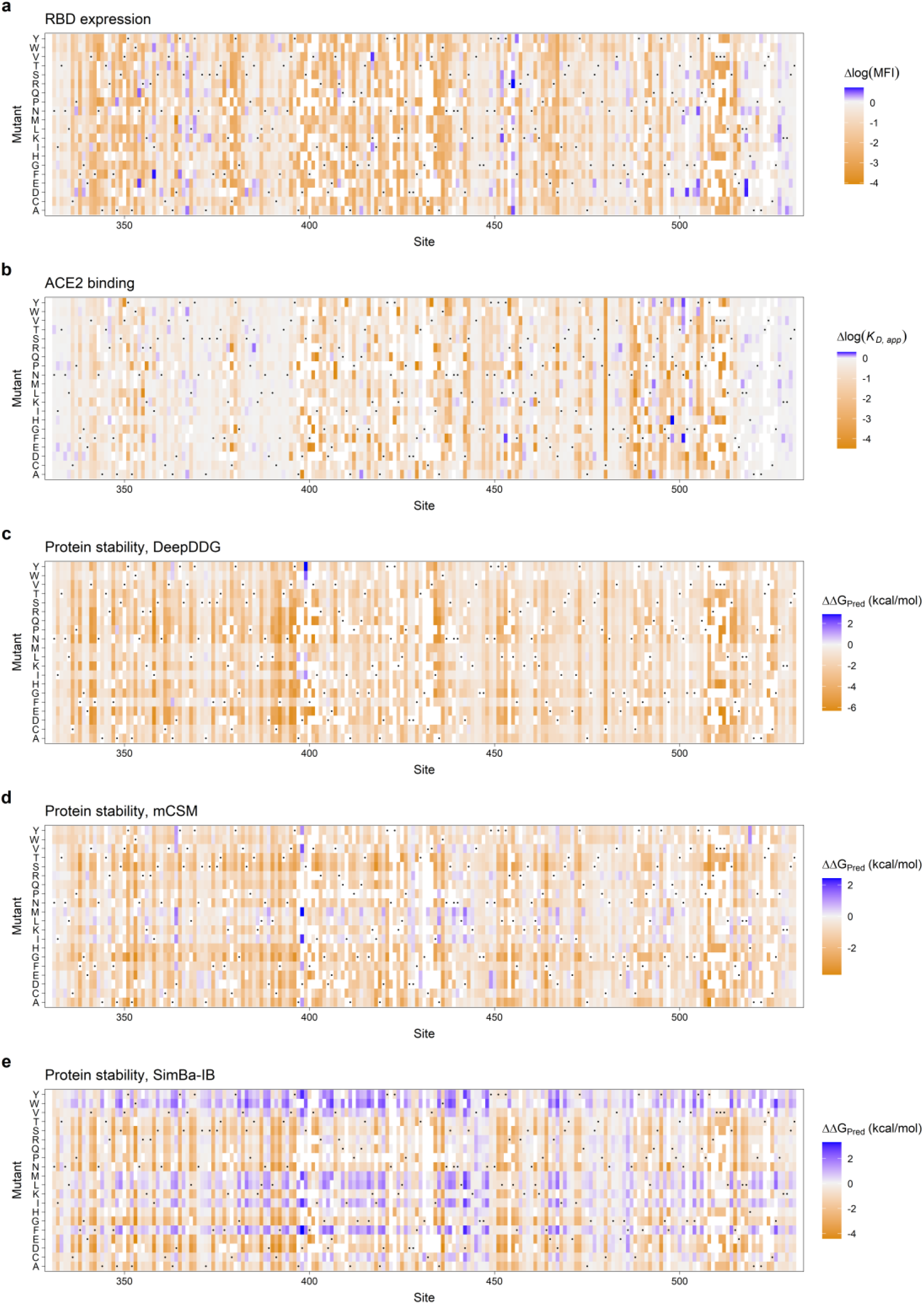
Heat maps of mutation effects on RBD experimental expression and ACE2 binding (from Starr et al.[53]) compared with computed stability change (this work). (**a**) expression; (**b**) ACE2-binding affinity. Protein stability change computed with: (**c**) deepDDG; (**d)** mCSM (**e**) SimBa-IB. Rectangles are colored by mutational effect according to scale bars on the right. Black dots indicate the wild-type residues. White rectangles represent mutations for which there is no data in the curated dataset.

The heat maps in **Figure 2** of experimental binding and expression do have some residual similarities, as also mentioned by Starr et al.[53] The computed stability effects provide estimates of the impact of all possible mutations in the RBD, and show some similarities to the expression data, notably with nearly neutral effects (grey) being common to both experimental and computed data in the N- and C-terminals and in some regions of the protein around 444-450 and a larger area around 470-488. In contrast, mutations in the region 388-396 have strong effects on protein stability but are nearly neutral with respect to expression and ACE2 binding. Overall, there is less similarity between ACE2 binding and the computed stability changes as perhaps expected.

### RBD sites with non-neutral effects cluster in the structured regions

**Figure 3** shows the effect of mutation at each site on expression, ACE2 binding and predicted protein stability mapped onto the RBD structure. The effect at each site is calculated as the average of the absolute effect (without sign) of all 19 mutations as a measure of the tolerance to mutation at each site. Mutations that affect expression are mainly located in the core RBD subdomain, and in particular in the central beta sheet and its flanking alpha helices (**Figure 3a**). Mutations that affect binding are located in the ACE2 binding subdomain or, similarly to mutations that affect expression, in the central beta sheet, while large parts of the RBD domain are tolerant to mutations with regard to ACE2 binding (**Figure 3b**). Mutations that affect protein stability are mainly located in the structured core of the RBD subdomain, similarly to the effect of mutations on expression (**Figure 3c-3e**). When evaluating the computational methods in this way, their ability to identify the most tolerant sites for mutation (or equally, the sites more likely to be neutral from an evolutionary perspective) becomes more evident than in the heat maps of **Figure 2**, while the better agreement with expression remains clear.

**Figure 3.**
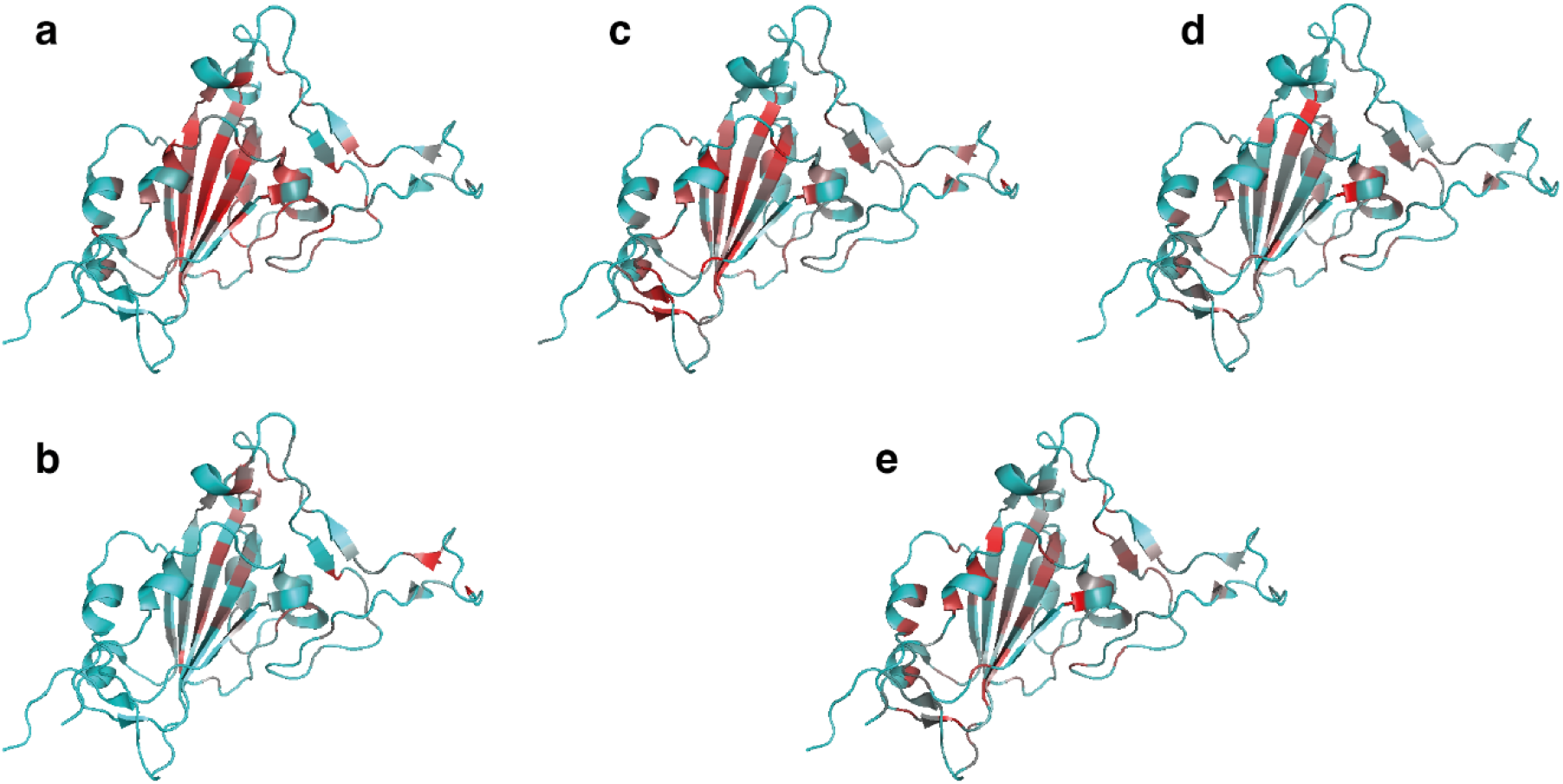
Effects of mutations mapped onto the SARS-CoV-2 RBD structure. Each residue is color-coded from cyan to red according to the average absolute effects of mutation. (**a**) RBD expression. (**b**) ACE2 binding. (**c**) Protein stability estimated by DeepDDG. (**d**) Protein stability estimated by mCSM, and (**e**) Protein stability estimated by SimBa-IB. Cyan indicates no effect, red indicates a strong effect. The color scales are relative within each panel. The ACE2 binding motif is shown on the right-hand side in each structure. (PDB: 6Z97, Chain A).

### Computed and experimental S-protein mutant properties correlate significantly

As discussed above, mutations in the RBD affect expression, ACE2 binding and protein stability differently, but with some overlap in site effects especially for protein stability and expression. To quantify this relationship, we plotted the predicted ΔΔG values for all mutations against the observed changes in RBD expression (**Figure 4a**) and ACE2 binding (**Figure 4b**), respectively, for each experimental structure used as input for the three methods (24 comparisons for all studied RBD mutations for expression and 24 comparisons for ACE2 binding). Correlations between RBD expression and ΔΔG for individual PDB structures are consistently observed, but their magnitude depends on the prediction method, with correlation coefficients ranging from 0.40 to 0.48 for DeepDDG, 0.27 to 0.34 for mCSM, and 0.12 to 0.22 for SimBa-IB (**Figure 4a**).

**Figure 4.**
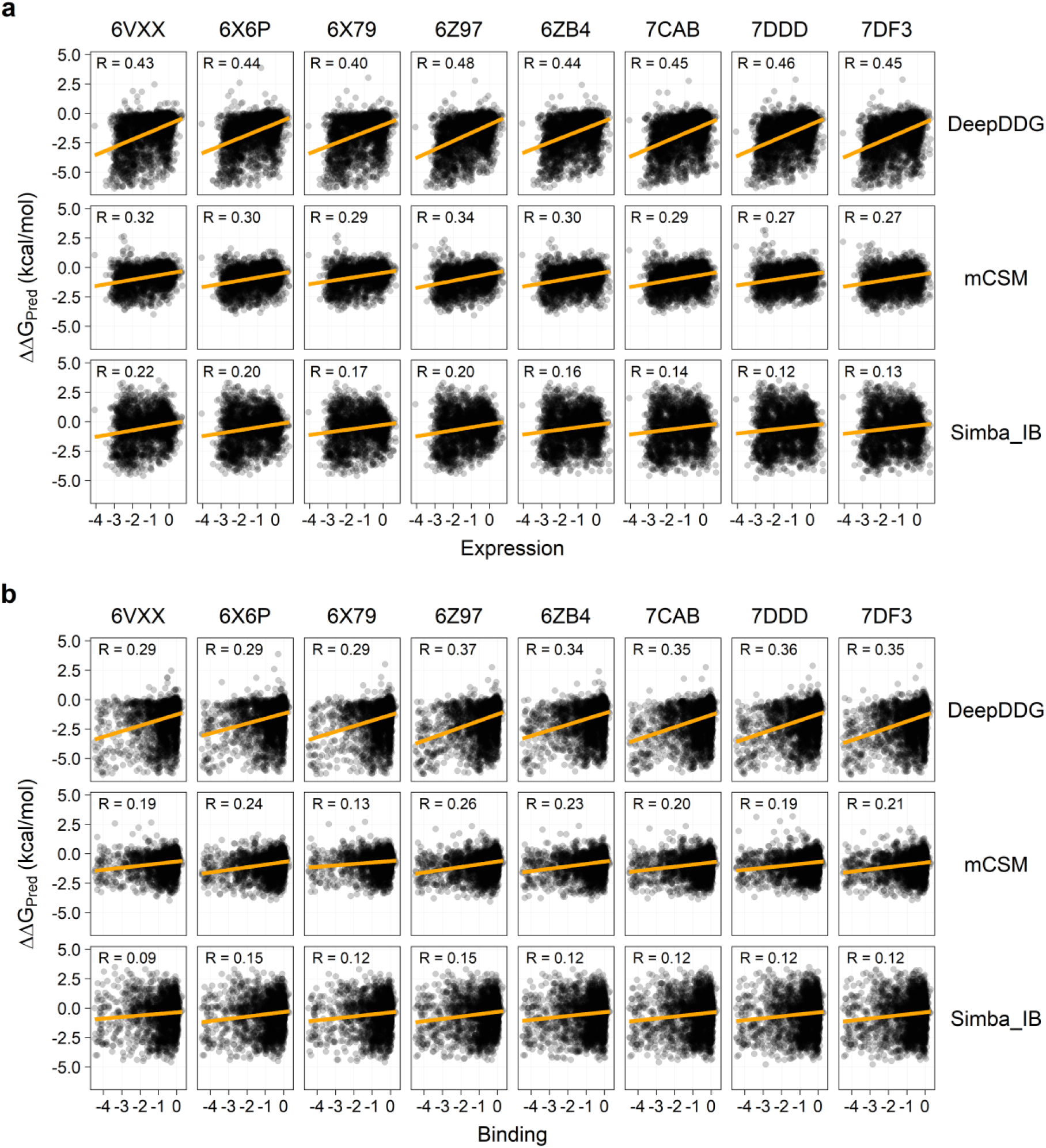
Predicted change in protein stability as a function of mutations effect on RBD expression and ACE2 binding. Three different prediction methods (DeepDDG, mCSM, and SimBa-IB; indicated on the right) were used to predict the change in protein stability for each mutation using eight different experimental structures of the S-protein (indicated on top). Orange lines indicate the resulting linear regression, and the correlation coefficients (R) are shown. The *p* values for all correlations are <0.001. **(a)** ΔΔG plotted against RBD expression, and **(b)** ΔΔG plotted against ACE2 binding.

The observed ACE2 binding and the predicted ΔΔG also correlate, depending on the prediction method (**Figure 4b**), which is in line with the correlation between experimental measures of expression and ACE2 binding (**Figure S3**). However, the correlations are weaker than for expression with correlation coefficients ranging from 0.29 to 0.37 for DeepDDG, 0.13 to 0.26 for mCSM, and 0.09 to 0.15 for SimBa-IB. In all 48 comparisons of computed and experimental data in **Figure 4**, the correlations are statistically significant at the 99% confidence level (p-values of linear regression < 0.01).

**Figure 5** shows the aggregate data for all structures, to account for structural heterogeneity effects. The relationships observed in **Figure 4** still hold true when averaging over all eight structures, i.e., our result is robust to structural heterogeneity. The effect of mutations on protein stability predicted using DeepDDG correlates better with both expression and ACE2 binding than using mCSM and especially SimBa-IB (**Figure 4** and **Figure 5**). Judging from **Figure 3c-e**, the three methods agree to a large extent on how they predict the tolerability of a site to mutation, whereas they differ in how they predict the effect of individual mutations (**Figure 2c-e**), and this is most likely the cause of the differences in the resulting correlations.

**Figure 5.**
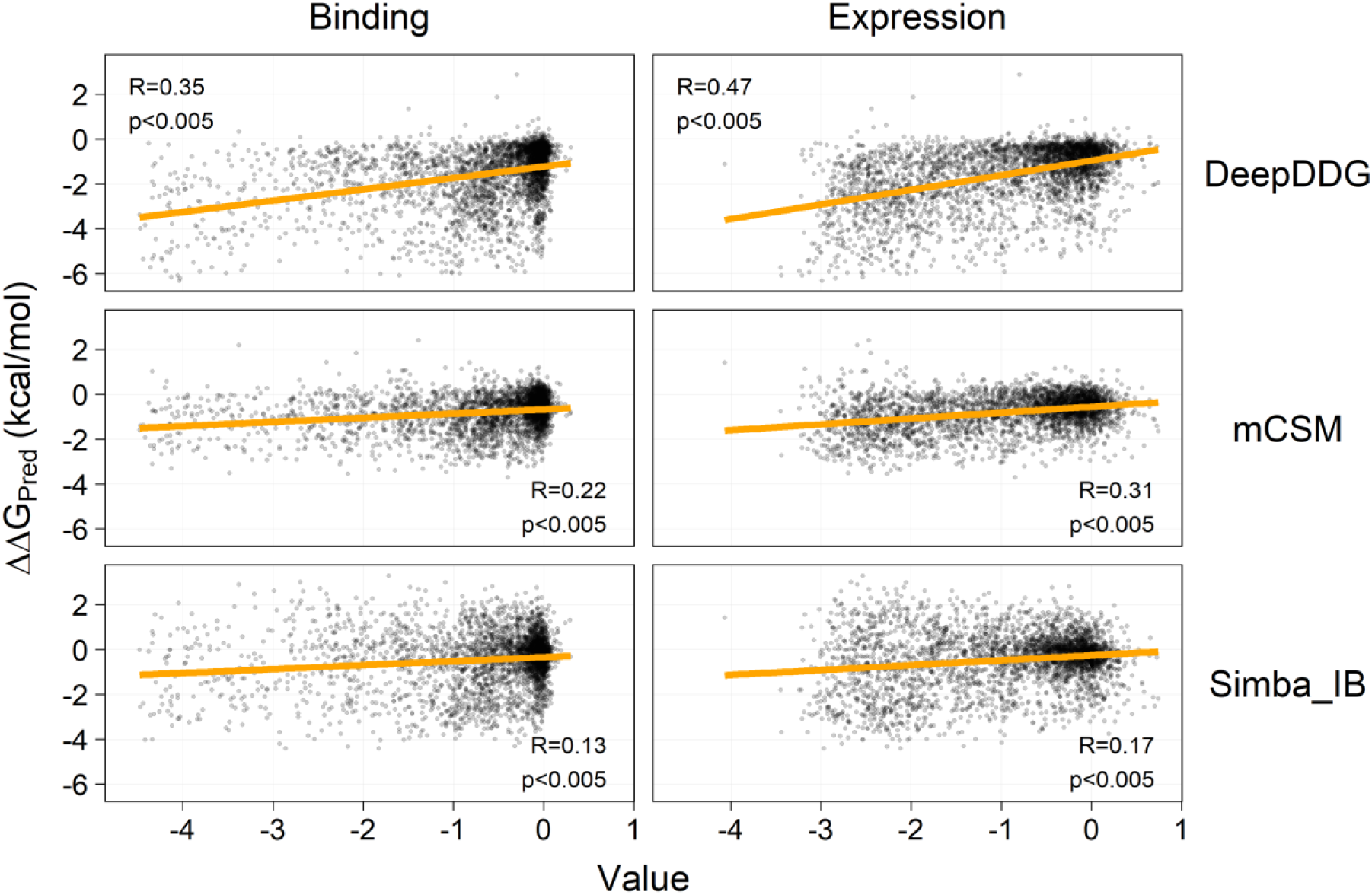
Average change in protein stability for the eight used structures vs. effect on RBD expression and ACE2 binding. Stability changes upon mutation for DeepDDG, mCSM, and SimBa-IB correlate with the experimental data at 99% significance (p-values of linear regression). Orange lines indicate the resulting linear regression.

Both the binding and expression effects are highly skewed, with an overrepresentation of data points between 0 and −1 (**Figure S2**). In order to adjust for this, we grouped the data into bins of 0.5-width and 0.25-width and calculated the mean expression and binding effect in each bin, and the mean predicted ΔΔG value for each bin. **Figure 6** shows these data as averages of all PDBs (data for individual structures in **Figures S4-S7**). The correlation increases substantially upon binning, with each data point more well-determined, whereas the *p*-values decrease substantially due to the few aggregate data points after averaging. Remarkably, the computational data correlate extremely well with the binned experimental data, especially for DeepDDG, much more than normally seen.[74] Considering that the models were developed to predict fold stability effects, not expression (which also depends on effects at the nucleic acid level) and that we used the full S-protein structures whereas the experiments express RBD on the yeast surface with expected modifications, this result is very surprising. One interpretation of this result is that broader functional properties of the mutant space of the S-protein are in fact largely determined by a few simple features, due to underlying correlations.

**Figure 6.**
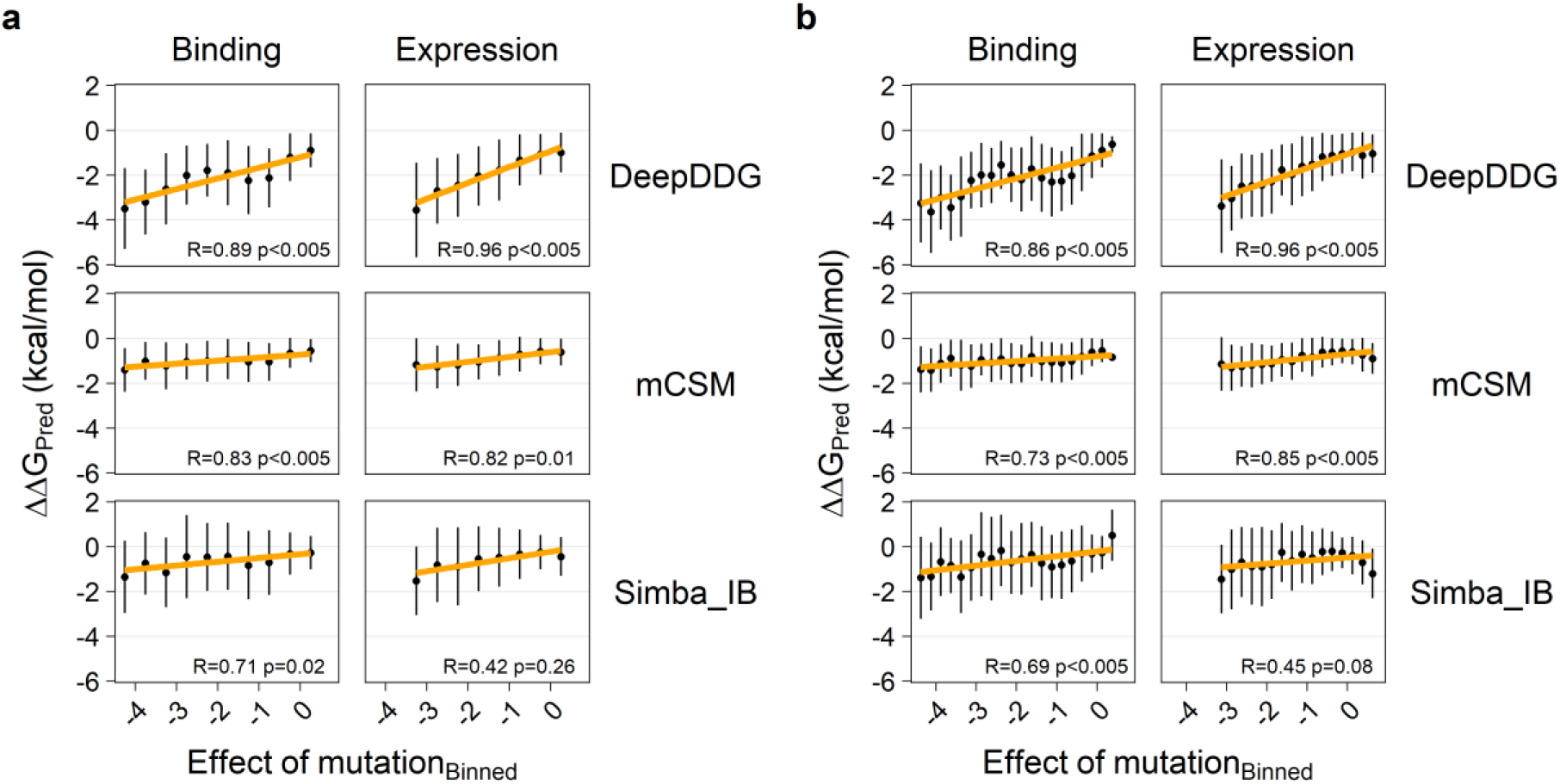
Predicted change in protein stability as a function of mutations effect on RBD expression and binding to ACE2 using binned data. Three different prediction methods (DeepDDG, mCSM, and SimBa-IB) were used to predict the change in protein stability for each mutation using eight different experimental structures of Spike protein. The average change in stability for the eight structures is used in the calculation. **(a)** Binding and expression data grouped in 0.5-value bins (**b**) Binding and expression data grouped in 0.25-value bins.

To understand the correlations on a per-site basis, **Figure 7** shows the relationships between the experimental and computed data averaged over sites, as an indicator of the site’s tolerance to mutations. As this removes some amino-acid specific variations between the three computer models used, the correlations now become more similar between the methods. In all cases, there is a significant correlation (99% confidence level, p-values of linear regression), such that sites more neutral to expression and ACE2 binding effects experimentally are also more neutral towards computed stability effects.

**Figure 7.**
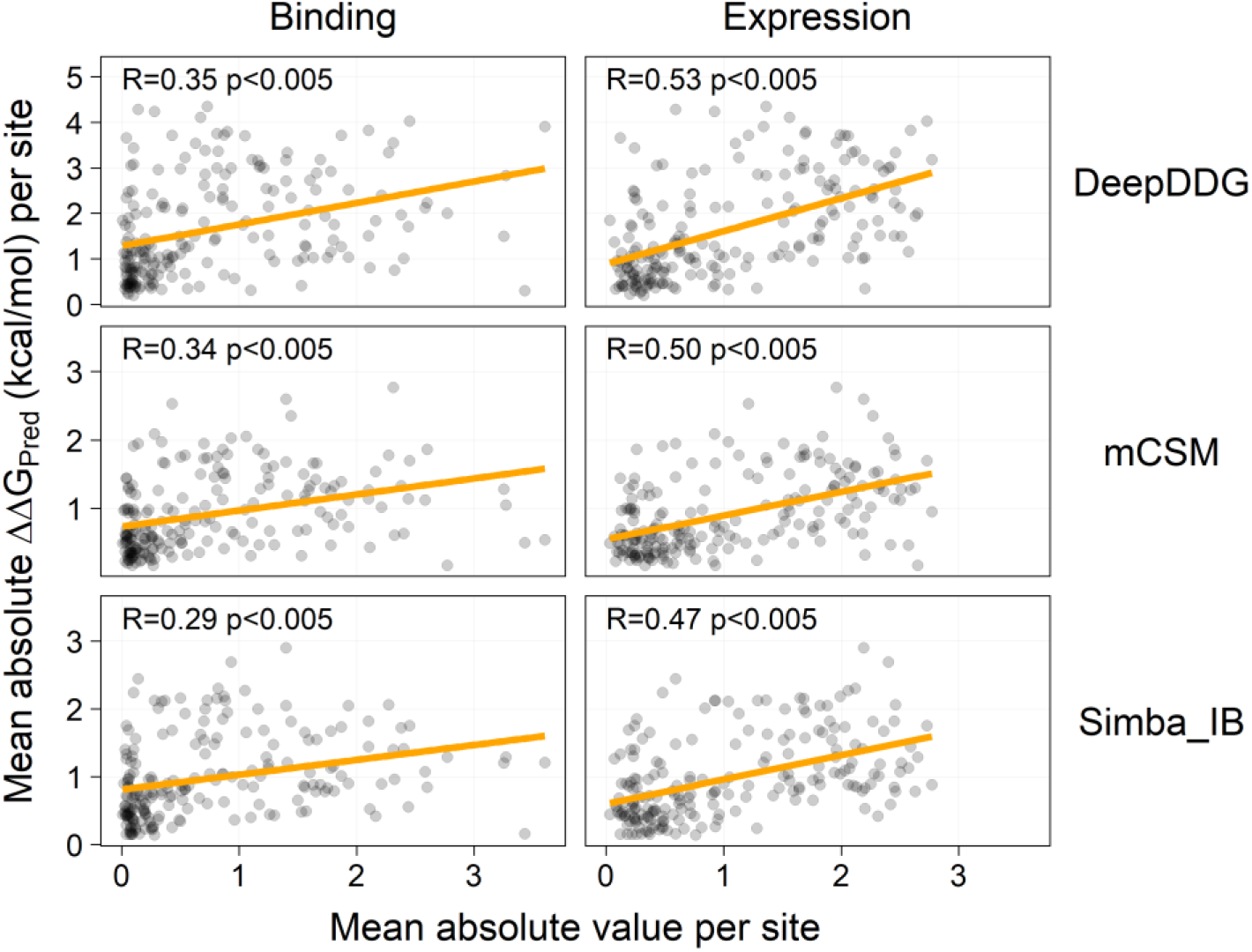
Site-averaged impact on ACE2 binding and expression vs. stability changes. This figure shows experimental-computational relationships for site-averaged properties (all mutations with data for each site averaged) as an indicator of the site’s tolerance to mutation.

Mutations in residues that are buried in the core of a protein tend to have larger effect on protein stability, which is also the case for the RBD domain, where mutations in the core subdomain are less tolerated (**Figure 3c**-**3e**) than mutations near the surface. To quantify if surface exposure of residues in RBD correlated with the effect of the mutations on expression and binding, we plotted these variables against the RSA for each site, and we see a moderate correlation (R = 0.29 for binding, and R = 0.43 for expression) with a notable overall tolerance to mutation at sites with high solvent exposure (**Figure S8**). As expected, there is also a good correlation between surface exposure and predicted stability changes, with correlation coefficients ranging from 0.27 (for SimBa-IB) to 0.60 (for DeepDDG; **Figure S9**).

### Concluding remarks and biological implications

The SARS-CoV-2 S-protein fusion with human ACE2 is a prerequisite for host cell entry.[9, 10] However, S-protein antibodies from previous infection or vaccines in the population bind the S-protein and neutralize some virus particles, thus reducing infectivity. During evolution of SARS-CoV-2, these two effects need to be balanced upon changes in the S-protein.[28] Most SARS-CoV-2 evolution until the omicron variant involved positive selection for better ACE2 binding[75], a feature maintained with omicron despite substantial evasion of existing antibodies via its many mutations in the S-protein.[76] In addition to these two, protein fold stability is commonly important during the evolution of new functionality, as it constrains the options available (function-stability tradeoffs)[29, 30, 77, 78] and has been speculated to play a role in SARS-CoV-2 S-protein evolution as well.[28] In order to understand and possibly predict SARS-CoV-2 evolution, an important health challenge,[28, 39] we evaluated such contributions of protein fold stability to the S-protein RBD mutation space.

Our work shows that computed protein stability effects correlate significantly for all 48 comparisons of data sets (8 structures, 3 methods, and two properties) with expression levels and to lesser extent ACE2 binding observed in experiments.[53] Thus, experimental mutant properties can partly be predicted, and the phenotypes may correlate, possibly due to underlying correlators between e.g. codon use, amino acid chemical properties, and site solvent exposure. Such correlations could impact the mutability of the SARS-CoV-2 S-protein and affect phenotype tradeoffs, and thus ultimately the virus evolution, given that S-protein fold stability in the prefusion state is an important property of SARS-CoV-2 mutants,[33] which suggests including protein stability into a fitness function of SARS-CoV-2.[28] To the extent that protein stability maintenance is an important selection pressure, it will affect the future SARS-CoV-2 S-protein evolution, including constraints on its antigenic drift.

Although single amino acid changes as analyzed above and in assays[79, 80] are likely to be not additive in real variants with multiple substitutions, due to amino acid correlation effects (within-protein epistasis), and possibly epistasis with other virus genes.[81–84] These epistasis effects haunt the protein evolution field and are not easily accounted for, but recent work suggests that substantial parts of the epistasis is already utilized,[85] although this remains to be studied in future work, and does not include potential inter-gene epistasis such as e.g. processing of S-protein RNA by non-structural proteins during virus replication within the host cell. Still, the maintenance of S-protein fold stability in the lipid surface of the virion is likely to be an important constraint on ACE2 binding and antigenic drift making the heat maps studied here of interest both as a proxy of expression and S-protein stability but possibly also as a model contribution to the computational estimates of the overall fitness function of SARS-CoV-2.

## Supporting Information

The supporting information pdf file contains additional data and analysis of relevance. The data set file Table_S1.csv contains the curated experimental data sets used for the study.

## Data availability statement

The full curated data used in this study are available as a csv file uploaded as supplementary information (Table_S1.csv), and scripts for analyzing these data are available at the web page https://github.com/ktbaek/Spike-ACE2.

## Acknowledgements

The Danish Council for Independent Research (Grant case: 8022-00041B) is gratefully acknowledged for supporting this work.

## SUPPLEMENTARY INFORMATION

**Figure S1.**
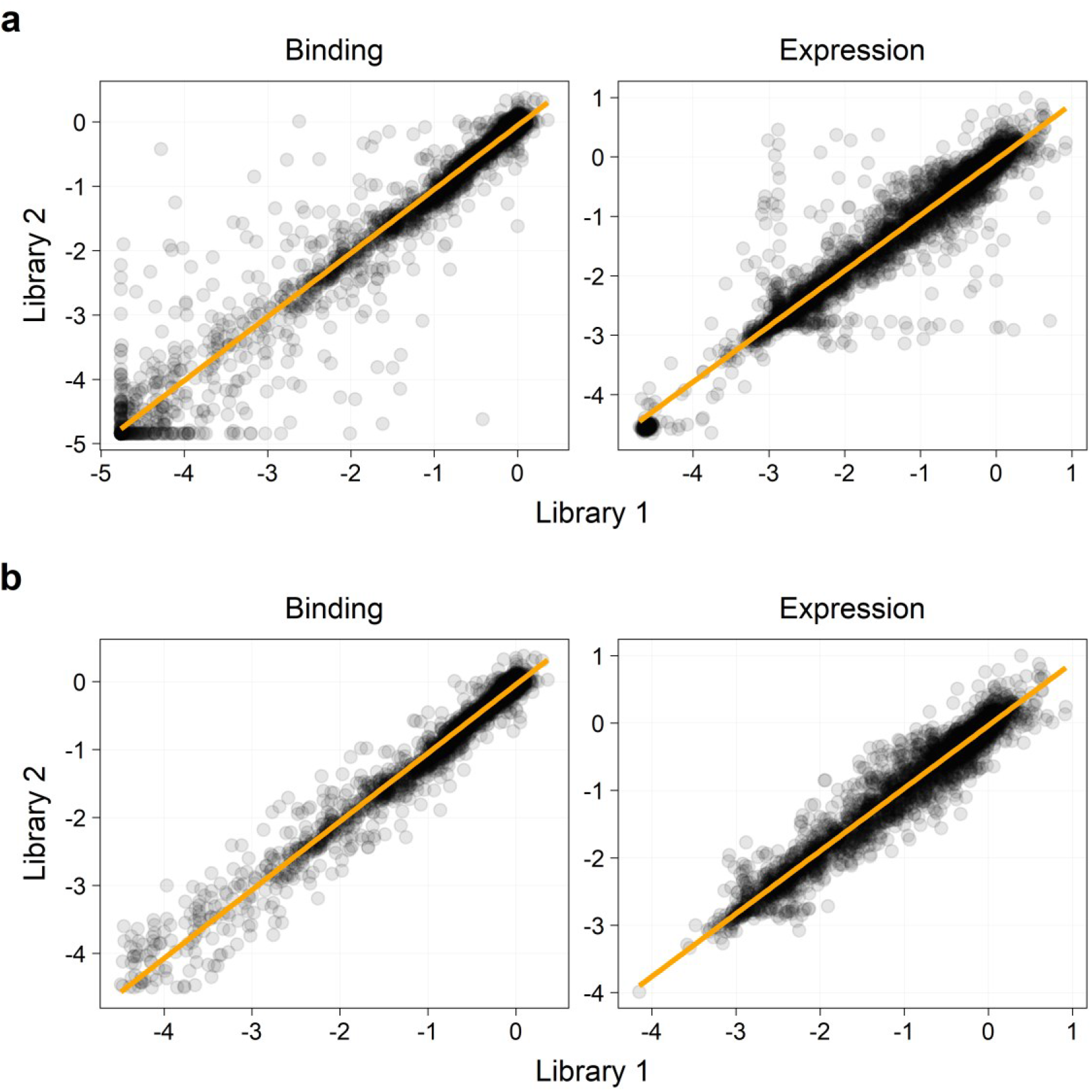
Correlation in effects on RBD expression and binding to ACE2 by two independent mutant libraries. **(a)** The original dataset from Starr et al.^1^ containing 4221 data points. **(b)** The dataset was curated by removing outliers (absolute residuals > 1), removing data points with binding less than −4.5 in one or both libraries, and removing stop codon mutations, resulting in a dataset of 3536 data points. The orange lines are the linear regression lines for the scatter plots.

**Figure S2.**
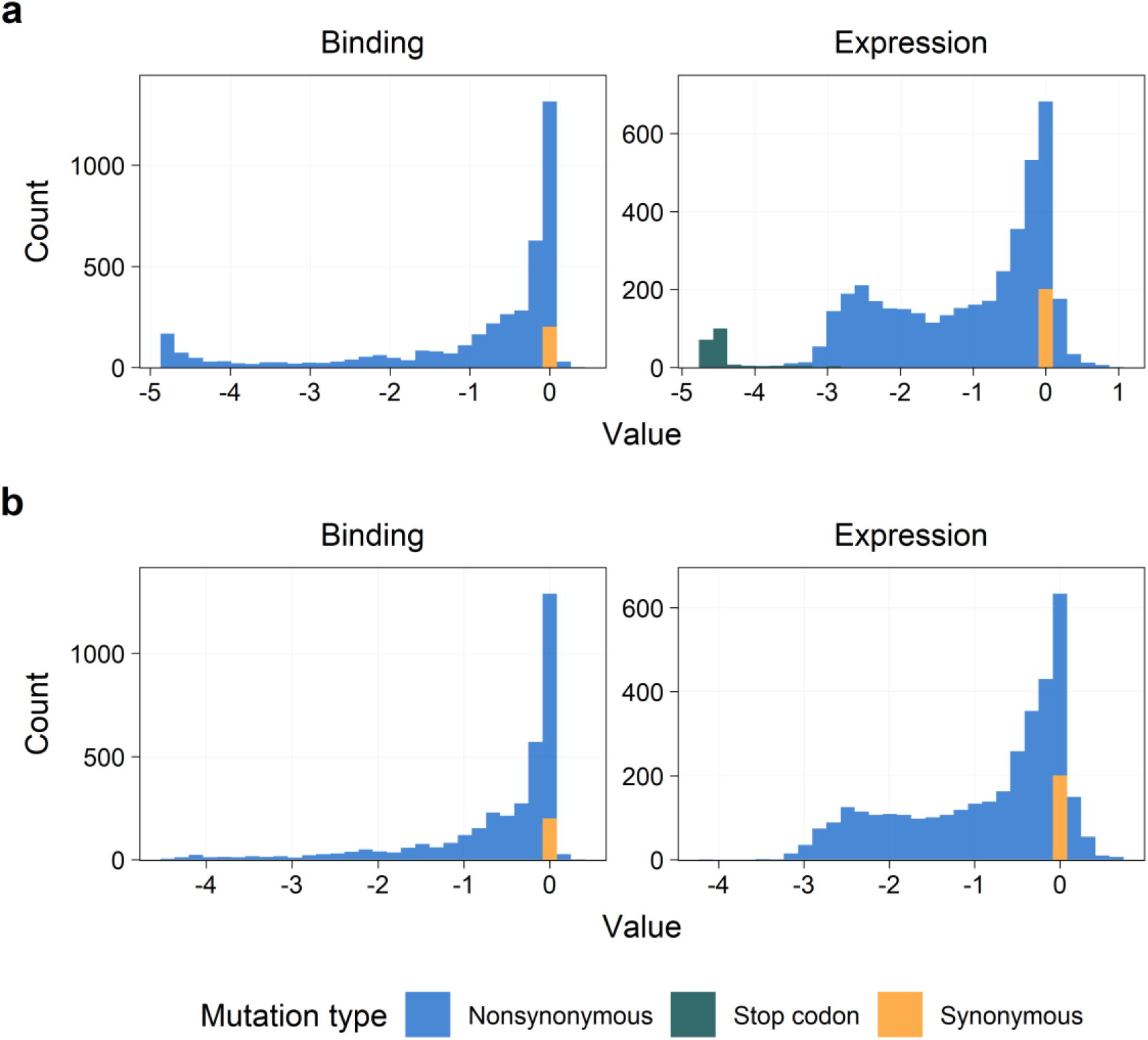
Distribution of mutations by effects on RBD expression and binding to ACE2. The distribution of mutations is shown for (**a**) the original dataset, and (**b**) the curated dataset. Color indicates the type of mutation. The average of the two libraries is used.

**Figure S3.**
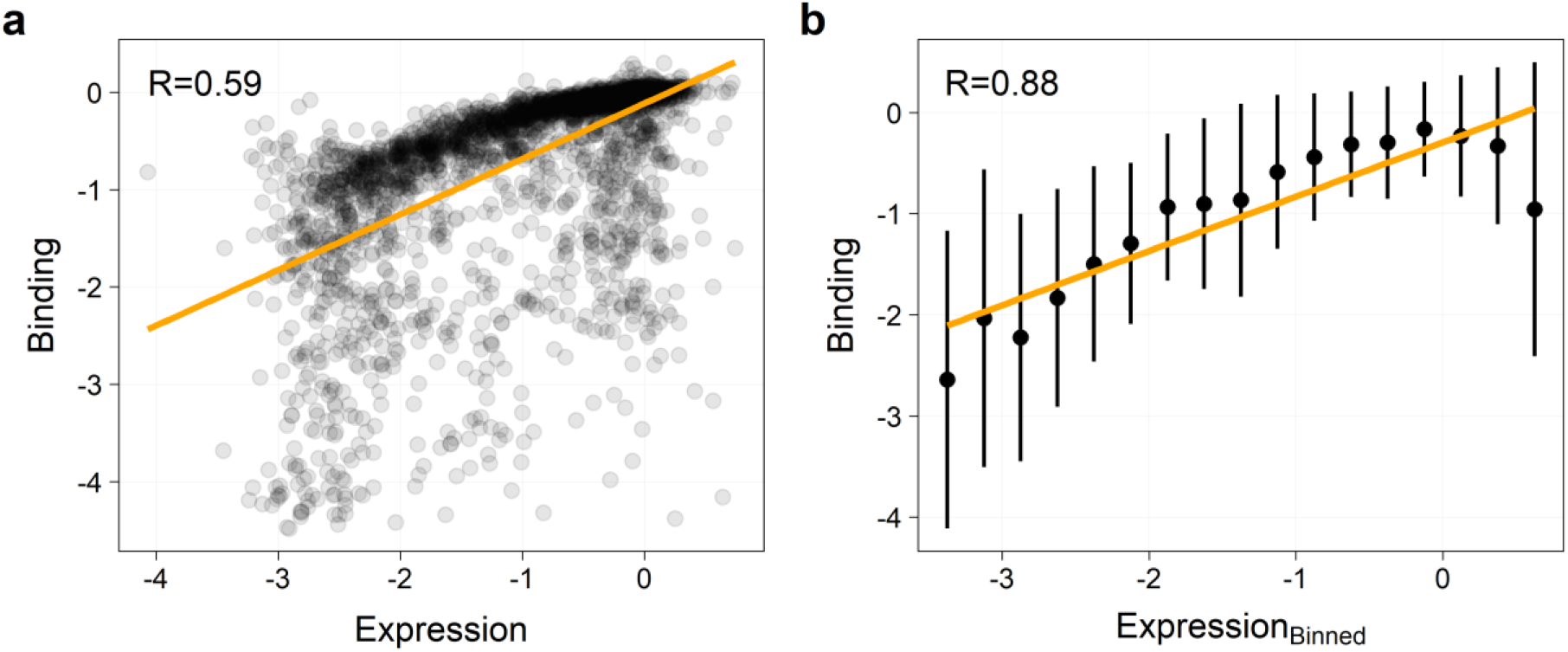
Relationship between mutation effect on RBD expression and binding to ACE2. **(a)** Each point represents one mutation’s effect on expression and binding. (**b**) Data points were binned according to the effect on expression in 0.25-wide bins, and points indicate the mean binding effect for each bin as a function of the middle expression effect in each bin. Vertical bars show the standard deviation in each bin. Orange line indicates the linear regression, and R is the correlation coefficient.

**Figure S4.**
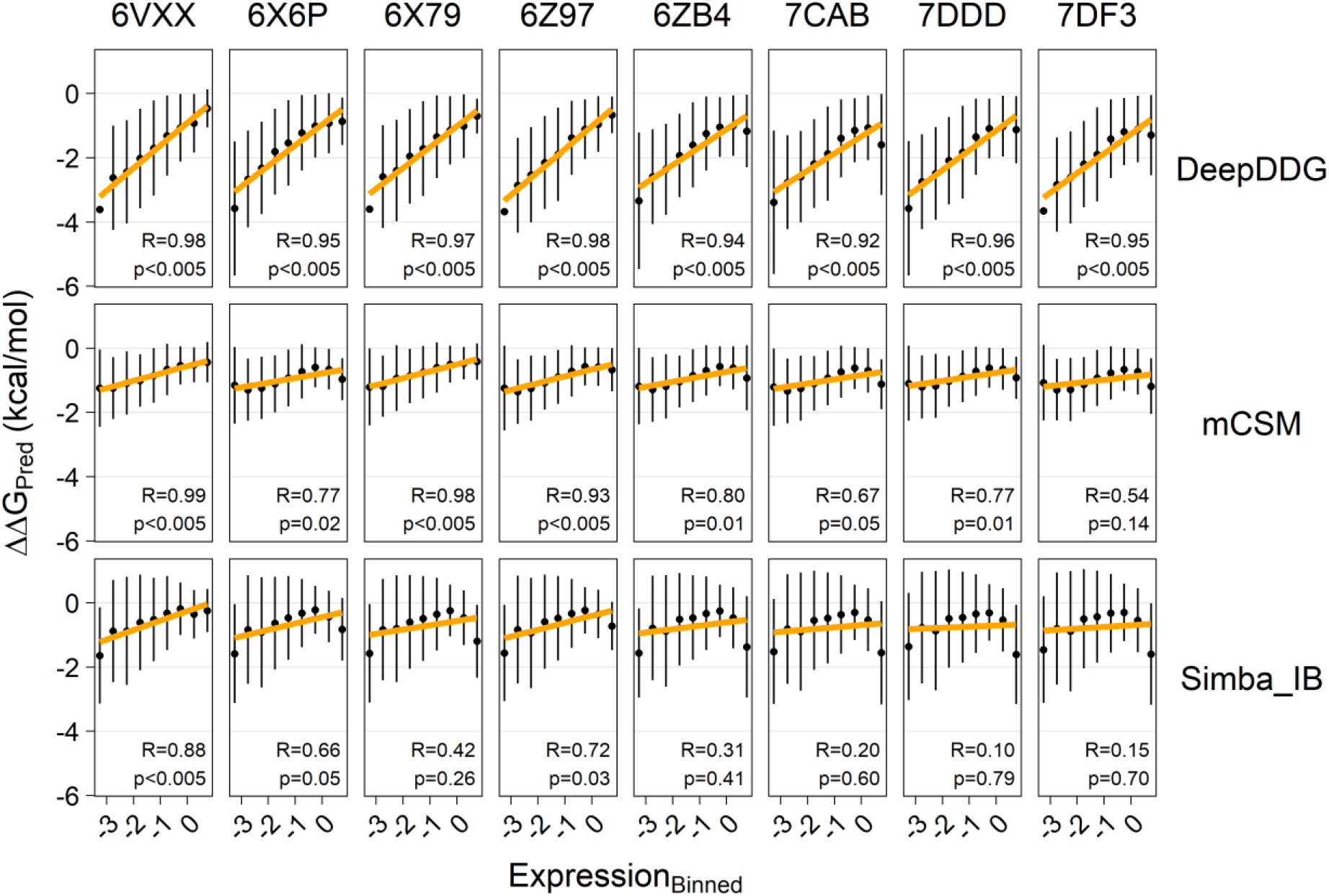
Predicted change in protein stability as a function of mutations effect on RBD expression – binned data (wide bins). Three different prediction methods (DeepDDG, mCSM, and SimBa-IB) were used to predict the change in protein stability for each mutation using eight different experimental structures of S-protein. Data points were binned according to effect on expression in 0.5-wide bins, and points show the mean ΔΔG for each bin as a function of middle expression effect in each bin. Vertical bars show the standard deviation in each bin. Orange lines indicate the resulting linear regression.

**Figure S5.**
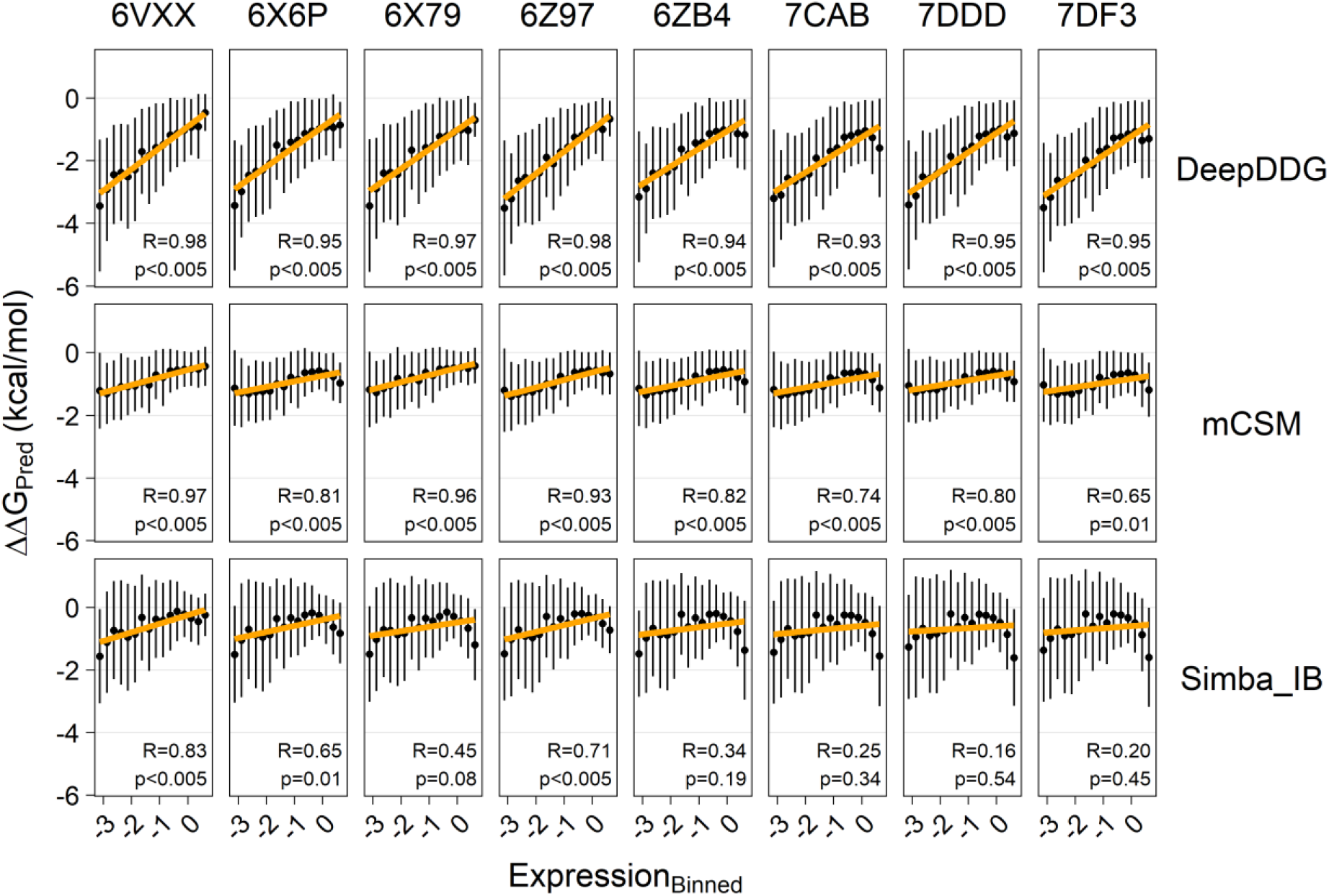
Predicted change in protein stability as a function of mutations effect on RBD expression – binned data (narrow bins). Three different prediction methods (DeepDDG, mCSM, and SimBa-IB) were used to predict the change in protein stability for each mutation using eight different experimental structures of S-protein. Data points were binned according to effect on expression in 0.25-wide bins, and points show the mean ΔΔG for each bin as a function of middle expression effect in each bin. Vertical bars show the standard deviation in each bin. Orange lines indicate the resulting linear regression.

**Figure S6.**
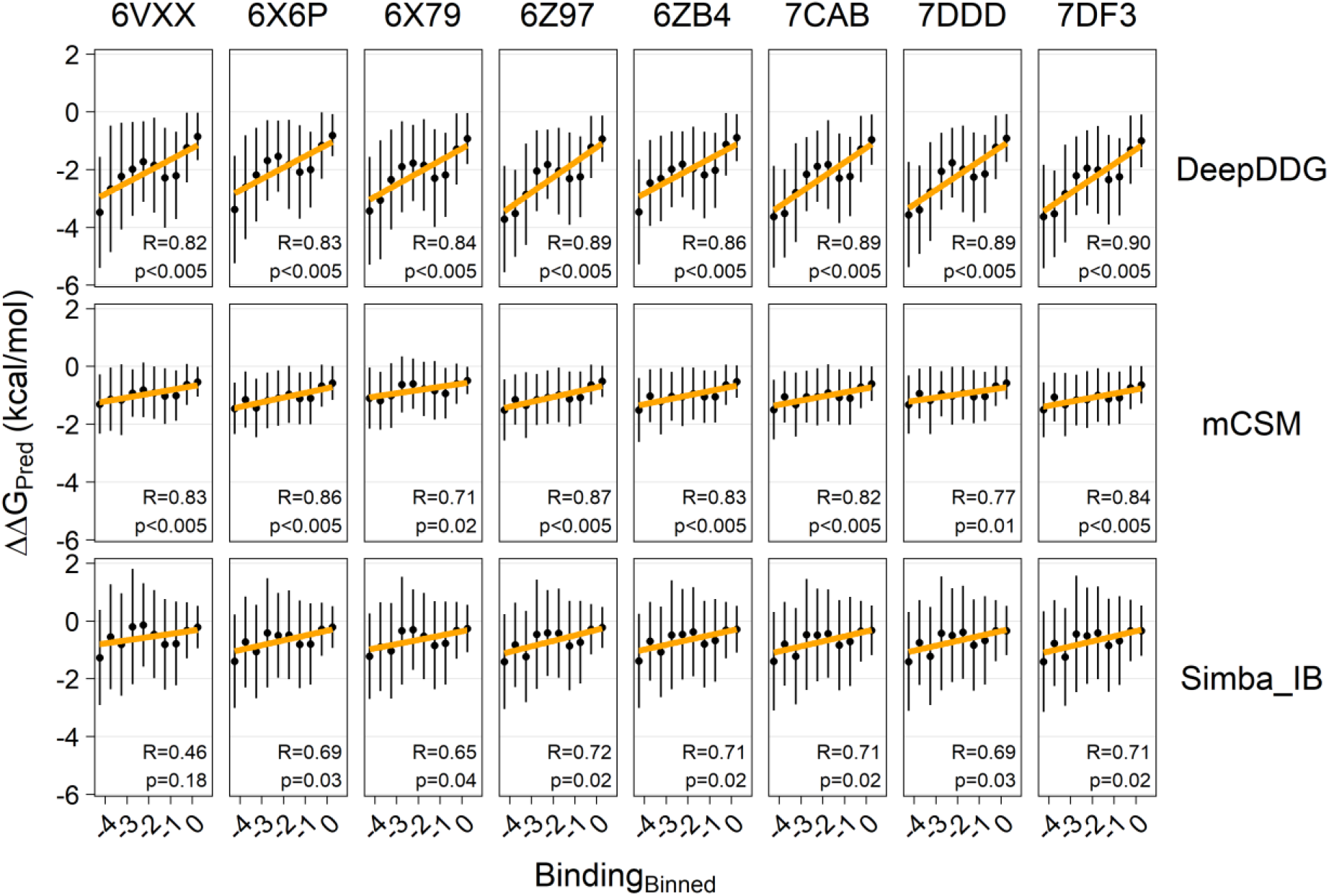
Predicted change in protein stability as a function of mutations effect on binding to ACE2 – binned data (wide bins). Three different prediction methods (DeepDDG, mCSM, and SimBa-IB) were used to predict the change in protein stability for each mutation using eight different experimental structures of S-protein. Data points were binned according to effect on binding in 0.5-effect bins, and points show the mean ΔΔG for each bin as a function of middle binding effect in each bin. Vertical bars show the standard deviation in each bin. Orange lines indicate the resulting linear regression.

**Figure S7.**
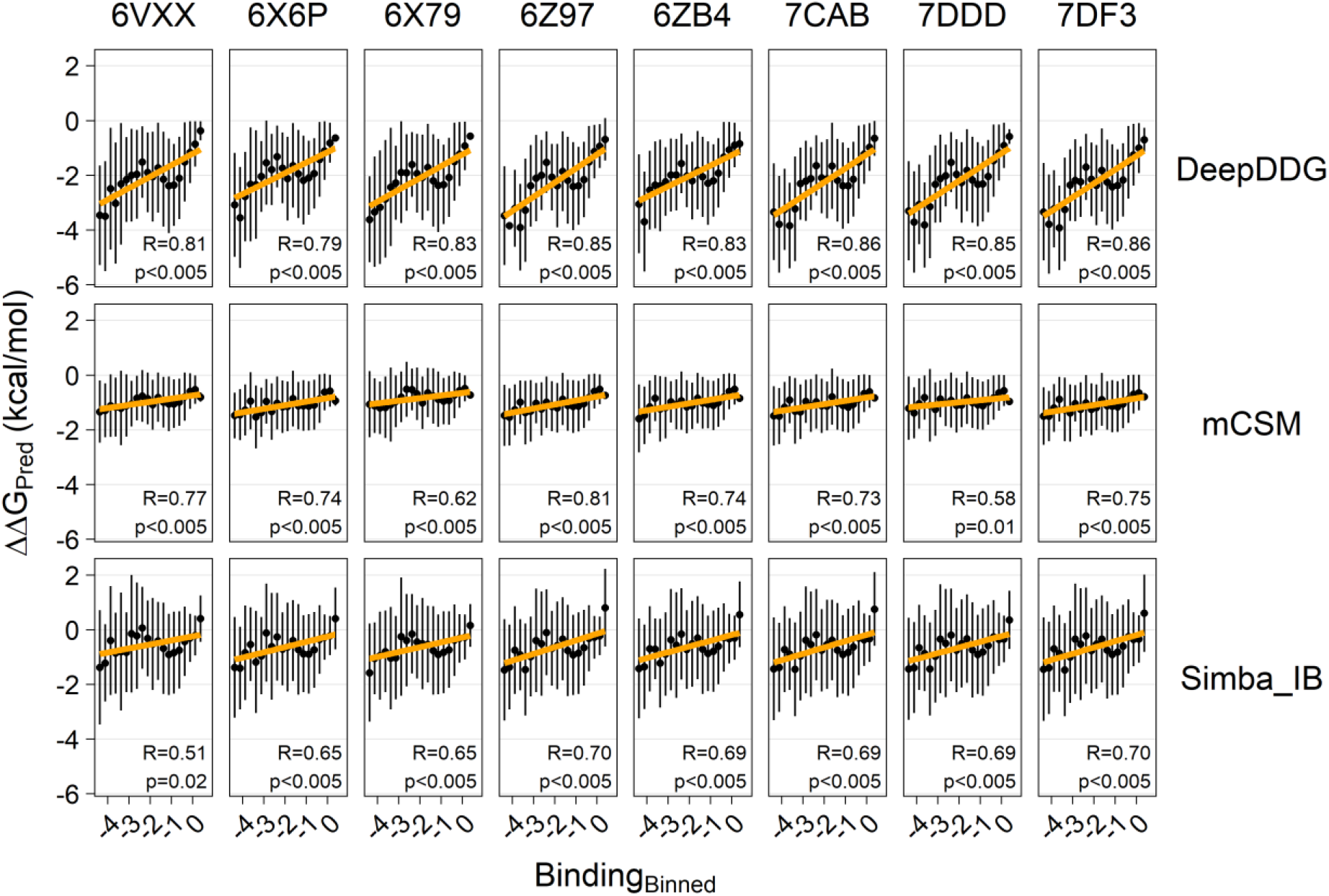
Predicted change in protein stability as a function of mutations effect on binding to ACE2 – binned data (narrow bins). Three different prediction methods (DeepDDG, mCSM, and SimBa-IB) were used to predict the change in protein stability for each mutation using eight different experimental structures of S-protein. Data points were binned according to effect on binding in 0.25-wide bins and points show the mean ΔΔG for each bin as a function of middle binding effect in each bin. Vertical bars show the standard deviation in each bin. Orange lines indicate the resulting linear regression.

**Figure S8.**
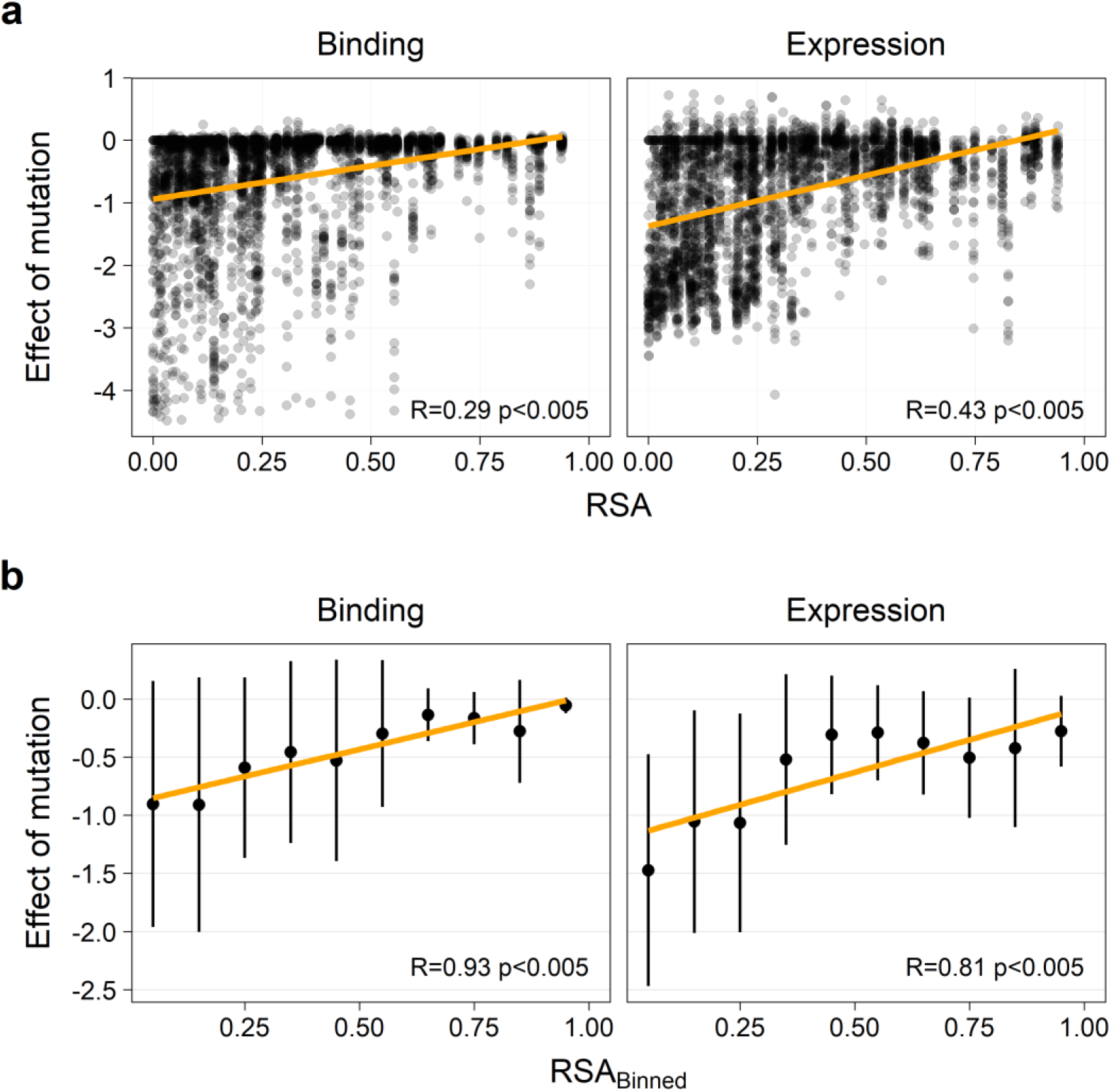
Relationship between RSA and the mutations’ effect on RBD expression and binding to ACE2. FreeSASA was used to calculate RSA for each wild-type residue using eight different experimental structures of S-protein. The average RSA value for the eight structures is used. (**a**) Each point represents one mutation. (**b**) Data points were binned according to RSA values in bins of 0.25 width, and points show the mean effect on binding or expression for each bin as a function of the middle RSA value in each bin. Vertical bars show the standard deviation in each bin. Orange lines indicate the resulting linear regression.

**Figure S9:**
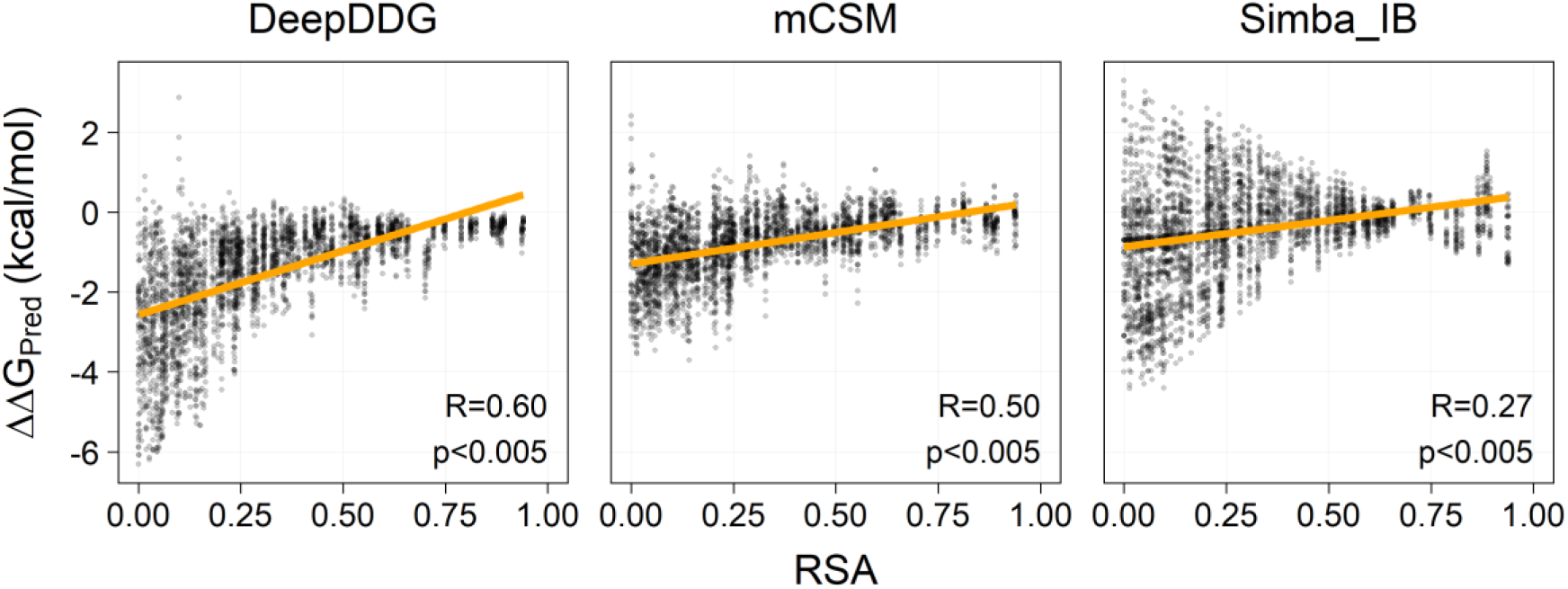
Change in stability as a function of relative solvent accessibility (RSA). Three different prediction methods (DeepDDG, mCSM, and SimBa-IB) were used to predict the change in protein stability for each mutation using the average of eight different experimental structures of S-protein. Each point represents one mutation.

